# Palaeoproteomic and genetic insights into millennial-scale dairy consumption in Armenia

**DOI:** 10.1101/2025.08.26.672348

**Authors:** Mariya Antonosyan, Anahit Hovhannisyan, Patrick Roberts, Lara M Cassidy, Satenik Mkrtchyan, Hasmik Simonyan, Arsen Bobokhyan, Levon Aghikyan, Ashot Piliposyan, Ruben Badalyan, Nzhdeh Yeranyan, Artak Gnuni, Hayk Avetisyan, Mikayel Badalyan, Armine Gabrielyan, Narineh Sarksian, Avetis Grigoryan, Hakob Simonyan, Meri Safaryan, Benik Vardanyan, Kristine Martirosyan-Olshansky, Zaruhi Khachatryan, Jana Ilgner, Levon Yepiskoposyan, Andrea Manica, Daniel G Bradley, Nicole Boivin, Shevan Wilkin

## Abstract

Dairy products are a key component of the diet in Armenia today, yet the origins of milk consumption, its historical development, and genetic adaptations related to milk digestion in the region remain understudied. Here, we investigate the co-evolution of dairying practices and lactase persistence in Armenia through combined palaeoproteomic and genetic analyses spanning from prehistory to the present. Our palaeoproteomic results provide direct evidence for milk consumption in prehistoric Armenia and, in conjunction with existing regional data, indicate that dairying was an established component of the South Caucasus subsistence economy from at least the late fourth millennium BCE. We found evidence of milk consumption from multiple species across archaeological periods, with cattle milk representing the earliest confirmed evidence from the Chalcolithic period. Notably, milk proteins were predominantly recovered from high-altitude sites, suggesting that dairy consumption served as an adaptive strategy for harsh highland environmental conditions. In contrast to the widespread evidence for milk consumption, our genetic findings indicate that the lactase persistence-associated variants never reached high frequencies throughout Armenian prehistory. This pattern suggests that the relatively low frequency of lactase persistence may reflect a predominant cultural adaptation to the fermentation of dairy products, which reduces lactose content and enables dairy use even in genetically lactose-intolerant populations.

## 1. Introduction

The incorporation of animal milk into human diets was a major innovation in prehistoric societies, enabling the supply of nutrients without slaughtering precious livestock. In marginal environments, milk additionally often functioned as a source of hydration and nutrients, supporting adaptive strategies in resource-scarce settings (Jeong et al. 2018; Tang et al. 2023). Moreover, the processing of milk into dairy products allowed for the production of food sources that could be stored and transported over long periods, sustaining mobile pastoral groups (Wilkin et al. 2020). These functional advantages of dairy products have been suggested to further boost long-term resilience and demographic success of groups that integrated dairy into their diets (Bocquet-Appel 2011; Wilkin et al. 2021; Ventresca Miller et al. 2022). Postweaning access to abundant sources of milk would not have been widely available to humans without the domestication of cattle, sheep, or goats. Zooarchaeological evidence, particularly those based on herd management and culling patterns, suggest that dairying may have emerged as early as the 8th millennium BCE (Vigne 2008; Vigne J-D 2007), shortly after the emergence of the first domesticated animals in the Near East (Vigne 2011). Meanwhile, the direct evidence for milk consumption detected by milk lipids as organic residues in archaeological pottery revealed that ruminants were exploited for their milk from the 7th millennium BCE onwards in Anatolia (Evershed et al. 2008; Nieuwenhuyse et al. 2015; Hendy et al. 2018), with this practice subsequently spreading across other regions at varying times (Dunne et al. 2018; Bleasdale et al. 2021; Debono Spiteri et al. 2016; Evershed et al. 2022; Outram et al. 2012; Wilkin et al. 2020; Jeong et al. 2018).

Located at the crossroads linking Mesopotamia and the Near East to the broader Eurasian continent, the South Caucasus region, including Armenia, served as a vital corridor for the exchange of people, knowledge and subsistence practices, including the spread of milk consumption northwards to the Pontic steppe by the 5th millennium BCE (Scott et al. 2022). This early tradition of milk use and processing is reflected in the enduring importance of dairy in the region. Today, dairy products are an essential part of daily diets in Armenia, with both fresh milk and processed dairy consumed regularly. Traditional processing follows three primary methods: additive coagulation (to produce cheese), fermentation (to produce yoghurt), or cream separation (to produce sour cream) (Hirata 2020). Among these, lactic acid fermentation is particularly widespread and central to local food culture, exemplified by *matsun*, Armenia’s traditional yoghurt (Afrikian 2012). In addition to being consumed as a fermented food, *matsun* also plays a versatile role in Armenian cuisine, serving as the base for tan, a refreshing summer drink, and as a key ingredient in *spas*, a traditional soup made with grains and yoghurt. When dried on bread (*choratan*), matsun can also be used as a starter culture for baking and when churned, it yields butter. Despite this rich dairy heritage, remarkably little is known about the origin and historical development of dairying practices in the region: a gap that palaeoproteomics analysis is uniquely positioned to address.

Adult milk consumption and lactose digestion are made possible by the ability to maintain postweaning production of the enzyme lactase–phlorizin hydrolase, which hydrolyses the milk sugar lactose into its component monosaccharides. The selection of adult lactase persistence (LP) phenotype and the consumption of prehistoric milk are suggested to be linked, reflecting coevolution between cultural innovations and genetic traits (Durham 1992; Tishkoff et al. 2007). LP is associated with at least five independently evolved SNPs, 13910C>T (rs4988235), −13907C>G (rs41525747), −13915T>G (rs41380347), −14009T>G (rs869051967), −14010 G > C (rs145946881)), located in the *MCM6* gene’s regulatory region approximately 14 kb upstream of the lactase *LCT* gene, with different variants predominating in different populations. As such, 13910C>T is particularly frequent in the populations of northern Europe and India, −13915T>G in the Near East, and other alleles in various East African and Sahelian pastoralist populations (Liebert et al. 2017; Ingram et al. 2009; Gallego Romero et al. 2012)). Nevertheless, fundamental questions regarding the origins of these alleles, their spatiotemporal selective dynamics, which seems to be a more complex process than previously thought (Evershed et al. 2022), as well as their prevalence in some of the undercharacterised regions, remain unanswered.

aDNA studies demonstrate the absence or low frequencies of the LP-alleles during the Neolithic period (Evershed et al. 2022; Ségurel & Bon 2017; Burger et al. 2007), suggesting that dairying practices developed largely in lactase non-persistent communities. This temporal mismatch between cultural and genetic evidence suggests that early dairy users likely relied on fermentation and other processing techniques to reduce lactose content and mitigate intolerance symptoms (Rosenstock et al. 2021). One of the earliest appearances of −13910C>T is reported in an Eneolithic Ukrainian individual, dated to 5,960 BP (Mathieson et al. 2018); (Segurel et al. 2020), though LP did not reach significant frequencies in Europe until after the Bronze Age (Mathieson et al. 2015). Although this increase has often been linked to the migration of populations from the Pontic-Caspian steppe around 5,000 years ago (Allentoft et al. 2015), this attribution is now debated (Burger et al. 2020). Recent findings highlight that many earlier studies relied on imputation to detect the allele, an approach now considered unreliable when ancestral haplotypes lacking the T allele for rs4988235 are present in ancient DNA samples (Burger et al. 2020; Ali et al. 2022). Moreover, other LP-associated alleles are underrepresented or absent in current reference panels, further limiting the accuracy of imputation. Additionally, contamination of ancient samples presents a significant challenge, as does the absence of stringent filtering protocols. This is particularly problematic given that the rs4988235 SNP involves a C>T transition, which is characteristic of ancient DNA damage patterns. Therefore, rigorous and direct screening approaches are essential for accurate inference of LP in ancient samples.

Here, we integrate proteomic and genetic analyses to investigate the history of milk consumption and lactose digestion in Armenia. We report evidence of dairy consumption in ancient populations through liquid chromatography-tandem mass spectrometry (LC-MS/MS) analysis of dental calculus from individuals dating from the Late Neolithic to the Middle Ages (Figure 1). In parallel, we present what is, to our knowledge, the first genetic screening of modern individuals from Armenia, and more broadly, in the Caucasus (Anguita-Ruiz et al. 2020), for SNPs associated with lactase persistence. We also screened all the available published ancient DNA dataset from the region for LP alleles. The Armenian population has demonstrated a high level of regional genetic continuity for well over 6,000 years (Hovhannisyan et al. 2025), making it a compelling population for tracing the temporal dynamics of this major cultural and nutritional adaptation through time. Thus, our integrative approach enables us to reconstruct patterns of milk use over time and evaluate the extent to which cultural and genetic adaptations to dairying co-evolved in this understudied region.

**Figure 1:**
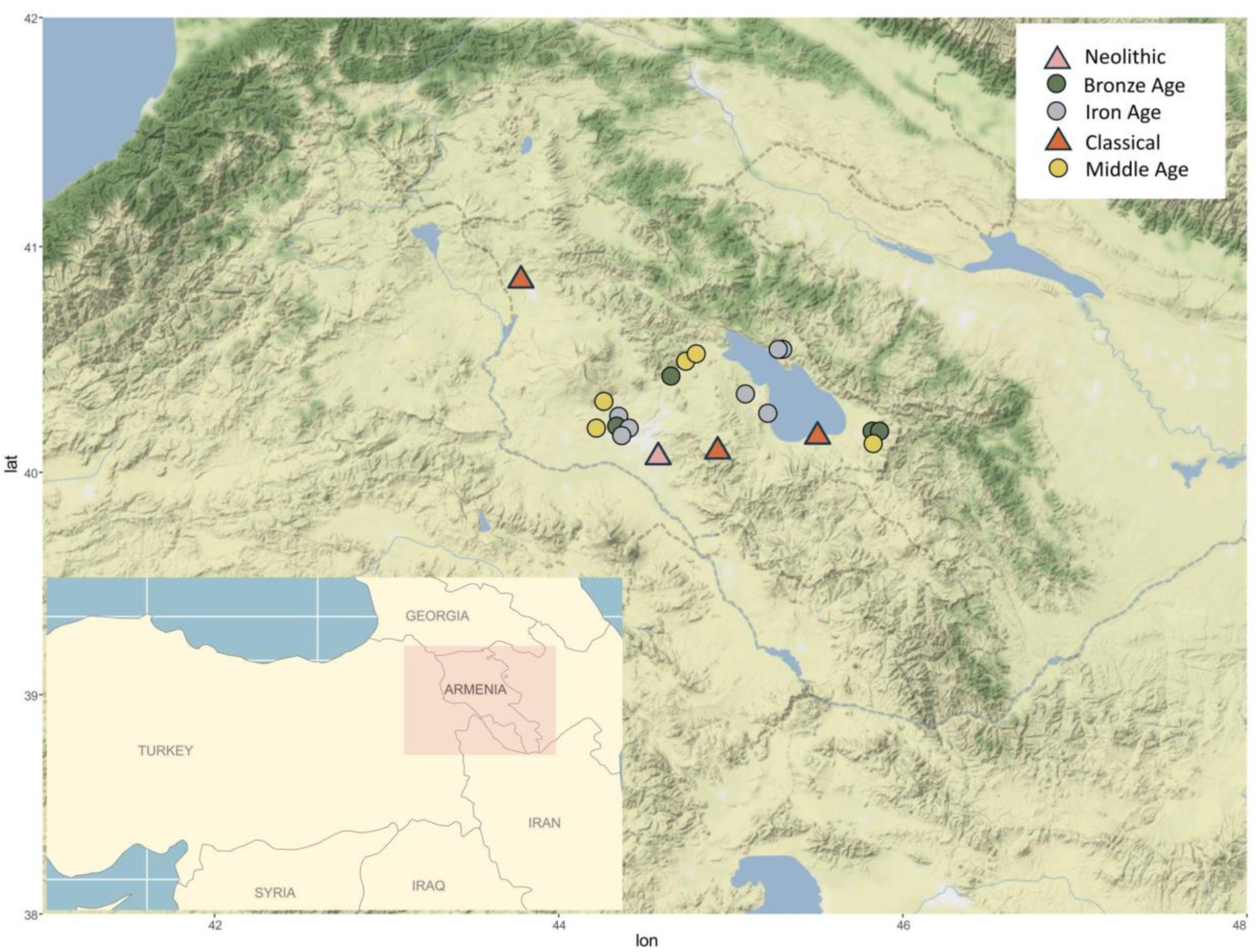
Locations of archaeological sites spanning from late Neolithic to Middle ages (site descriptions are presented in supplemental text)

## 2. Materials and Methods

### 2.1. Protein analysis

#### 2.1.1. Sample collection

To examine ancient dairy consumption history, we analysed ancient proteins extracted from the dental calculus of 76 individuals excavated from 22 archaeological sites across Armenia spanning the Late Neolithic to Medieval ages (Figure 1). The detailed archaeological contexts for each site are described in the Supplementary text. The list of individuals, including their age and sex profiles, is presented in Supplementary Table 1.

Dental calculus samples were collected in a dedicated clean laboratory at the Institute of Molecular Biology, NAS Armenia, under controlled conditions to reduce the risk of contamination. Nitrile gloves, surgical masks, and hair nets were worn throughout the sampling process to minimise the risk of modern contamination. Calculus was carefully detached from teeth using dental scalpels. To prevent cross-contamination between individuals, scalpels, gloves, and work surfaces were thoroughly cleaned with 75% ethanol between each sampling. The calculus samples were immediately transferred into sterile 1.5 ml Eppendorf tubes and stored until protein extraction, carried out at the dedicated Palaeoproteomics Laboratory at the Max Planck Institute for Geoanthropology (formerly MPI for the Science of Human History). Before extraction, 5-10 mg of calculus was weighed out in a microcentrifuge tube, and weights were recorded (see Supplementary Table 1). Additionally, 2-3 mg of ancient sheep bone powder was prepared as a positive control.

#### 2.1.2. Protein extraction

Samples were extracted using a single-pot, solid-phase enhanced sample preparation (SP3) modified for archaeological dental calculus samples (Wilkin et al. 2020; Hughes et al. 2019; Hughes et al. 2014; Cleland 2018). The full protocol can be found on protocols.io, DOI: https://dx.doi.org/10.17504/protocols.io.bfgrjjv6. Samples (including the positive control and extraction blank) were demineralised in 500 µl of 0.5 M Ethylenediaminetetraacetic acid (EDTA) for 4-7 days at room temperature. Following demineralisation, samples were centrifuged at 20,000 rcf for 10 min, and 400 μl of the supernatant was removed and retained. The remaining samples were denatured with 200 μl of 6 M guanidine hydrochloride (GuHCl), as well as reduced and alkylated 30 μl of 100 mM alkylated with chloroacetamide (CAA) / 100 mM Tris(2-carboxyethyl)phosphine (TCEP). Samples were then incubated at 99 °C for 10 min. Afterwards, 20 μl of a 20 μg μl^−1^ 50/50 mixture of hydrophilic and hydrophobic SeraMag SpeedBeads was added to each sample. To increase protein– bead adhesion, 350 μl of 100% ethanol was then added to each tube. Samples were then placed on the Thermoshaker at 1,000 rpm, 24 °C for 5 minutes. Afterwards, tubes were placed on a magnetic rack, which induced the migration of beads (carrying the proteins) to the wall of the tube. The sample/ethanol solution was removed and retained. The beads were then washed with 200 μl 80% ethanol. Upon washing, 100 μl of 100 mM ammonium bicarbonate was added to each tube, as well as 0.2 μg of trypsin to initiate enzymatic digestion. The samples were incubated on a Thermoshaker at 750 rpm, 37 °C for 10 minutes. After, they were resuspended and left to incubate overnight (approximately 18 hours). Following digestion, the tubes were centrifuged at 20,000 rcf for 1 minute and returned to the magnetic rack. The supernatant was transferred to clean tubes. To stop trypsin digestion, each sample was acidified using 5% trifluoroacetic acid (TFA). The acidified samples were then centrifuged at maximum speed for 5 minutes to pellet any residual non-proteinaceous material. Peptides were then desalted with C18 StageTips. Samples were sent on stage tips to the Functional Genomics Center Zürich, ETH/University of Zürich for LC-MS/MS analysis.

#### 2.1.3. LC–MS/MS analysis

LC-MS/MS analysis was performed on a Q Exactive mass spectrometer (Thermo Scientific) equipped with a Digital PicoView source (New Objective) and coupled to a nanoAcquity UPLC (Waters). The solvent composition at the two channels was 0.1% formic acid for channel A and 0.1% formic acid, 99.9% ACN for channel B. The column temperature was maintained at 50 °C. For each sample, 4 μl of peptides were loaded on a commercial ACQUITY UPLC M-Class Symmetry C18 Trap column (100 Å, 5 μm, 180 μm × 20 mm, Waters) followed by ACQUITY UPLC M-Class HSS T3 column (100 Å, 1.8 μm, 75 μm × 250 mm, Waters). The peptides were eluted at a flow rate of 300 nl min−1 by a gradient from 8 to 22% B in 49 min and to 32% B in an additional 11 min. The column was subsequently washed by increasing to 95% B and holding 95% B for 5 minutes before re-establishing the initial conditions.

The mass spectrometer was operated in data-dependent mode, acquiring a full-scan mass spectra (300-1,700 m/z) at a resolution of 70,000 at 200 m/z after accumulation to a target value of 3,000,000 and a maximum injection time of 110 ms followed by HCD fragmentation on the 12 most intense signals per cycle. HCD spectra were acquired at a resolution of 35,000 using a normalized collision energy of 25 and a maximum injection time of 110 ms. The automatic gain control was set to 50,000 ions. Charge state screening was enabled. Singly, unassigned and charge states higher than seven were rejected. Only precursors with intensity above 9,100 were selected for MS/MS measurement. Precursor masses previously selected for MS/MS measurement were excluded from further selection for 30 s and the exclusion window was set at 10 ppm. The samples were acquired using internal lock mass calibration on m/z 371.1012 and 445.1200.

#### 2.1.4. Data analysis

Generated raw LC-MS/MS spectra were converted to Mascot generic files using MSConvert from the ProteoWizard software package v.3.0.11781 by selecting the top 100 peaks from each spectrum. Each sample was searched using Mascot (Matrix Science; v.2.6.0) against the SwissProt database in combination with a custom curated dairy database consisting of 245 dairy livestock milk protein sequences (Wilkin et al. 2020). Mascot searches were conducted with a peptide mass tolerance of 10 ppm, with fragment mass tolerance of 0.01 Da and an allowance for monoisotopic mass shifts. Up to three missed cleavages were allowed. Carbamidomethyl of cystine (C) was set as a fixed modification, together with deamidation of asparagine (N) and glutamine (Q) and oxidation of methionine (M) as variable modifications.

To estimate the validity of peptide identifications and summarise the findings, the results were processed through an internally created tool, MS-MARGE (Hagan 2018), available for use via Bitbucket: https://bitbucket.org/rwhagan/ms-marge/src/master/. As input, MS-MARGE accepts a csv file exported from a Mascot MS/MS ion search against an amino acid database containing decoyed sequences with the Group Protein Families option turned off. MS-MARGE calculates the false discovery rate (FDR) at the PSM and protein level by comparing the number of forward sequence hits to the number of reverse sequence hits. For each sample, the protein false-discovery rate was targeted at under 5% and a peptide false-discovery rate at under 2%. A minimum of two individual peptide spectral matches were required for each specific protein identification, and only peptide spectral matches with an e-value of below 0.01 were accepted. To further investigate the taxonomic assignment, BLAST and Uniprot searches were performed on the identified peptides.

#### 2.1.5. Proteome preservation assessment

Proteome preservation of each dental calculus sample was assessed using a curated Oral Signature Screening Database (OSSD) (Bleasdale et al. 2021; Wilkin et al. 2021; Ventresca Miller et al. 2023). The OSSD contains common contamination proteins, including those encountered in the environment and laboratory, common bacteria found in the human microbiome, and human immune proteins that are regularly identified in the oral cavity. The screening approach is based on the premise that well-preserved ancient dental calculus retains a representative profile of oral microbiota and immune proteins typically present in saliva. Consequently, calculus samples exhibiting a strong oral proteomic signature—defined by meeting the OSSD threshold—are considered more likely to yield authentic insights into ancient dietary practices. To meet the OSSD criteria, a sample must contain a minimum of 15 identified OSSD proteins, with over 50% attributed to oral bacterial taxa and/or human inflammatory response proteins. Sample authenticity is further supported by the absence of dietary proteins in all positive (archaeological sheep bone with known proteome) and negative controls (extraction blank). Supplementary Table 2 contains the overall count of OSSD proteins pulled from our filtered results, as well as the result of the oral microbiome protein identifiers.

### 2.2. Genetic analyses

#### 2.2.1. Modern data

We screened five lactase persistence-associated variants in the full genomes of 36 Armenians, each with all four grandparents (4GP) originating from the same region within the Armenian highlands (Hovhannisyan et al. 2025; Mallick et al. 2016). In addition, we screened genotype data from 27 4GP Armenian individuals (Hovhannisyan et al. 2025) and 99 non-4GP Armenians for the −13910C>T allele (Haber et al. 2016). The data has been processed as it is described in Hovhannisyan et al. 2025 (Hovhannisyan et al. 2025). plink2 was used to convert vcf file to the PLINK binary format, calculate allele frequency and Hardy-Weinberg equilibrium for the given locus (Purcell et al. 2007).

#### 2.2.2. Ancient data

We downloaded sequencing data for all previously published ancient samples from Armenia (Lazaridis et al. 2022; Allentoft et al. 2015; Antonio et al. 2024; Damgaard et al. 2018; Wang et al. 2019; Bobokhyan et al. 2024). When available, the data were downloaded and aligned from a fastq file or, alternatively, realigned from a bam file. Adapter sequences were removed using Cutadapt (Martin 2011) and AdapterRemoval (Lindgreen 2012) for samples sequenced with single-end and paired-end technologies, respectively. The genomes were aligned to GRCh37 with decoy contigs (hs37d5) human reference genome using BWA software (Li 2013) with non-default parameters -l 16500, -n 0.02 and - o 2. Aligned reads were filtered for a mapping quality filter of 25 using samtools version 1.7 (Danecek et al. 2021) and for a minimum read length equal to 34 bp. For manual variant screening, we used GATK pileup –verbose (McKenna et al. 2010), filtered for base quality 30, mapping quality 25 and read length 34 bp. The two high-coverage Bronze Age individuals (Bobokhyan et al. 2024) were genotyped with HaplotypeCaller in GATK. mapDamage was used to rescale base qualities (Jónsson et al. 2013).

#### 2.2.3. Imputation

Biallelic variants and genotype likelihoods were called with “-mpileup” and “-call” in bcftools v1.1 (Danecek et al. 2021), respectively. In order to detect the haplotypes on which the LP allele arose or which potentially carry the allele, we imputed both captured and shotgun ancient genomes with GLIMPSE v1.0.1 using 1000 Genomes Project haplotype reference (1000 Genomes Project Consortium 2015). To avoid batch effect, imputation was conducted with the samples independently. Genomes were initially divided into 2 Mb windows with a 200 kb buffer using GLIMPSE_chunk (Rubinacci et al. 2023), then imputed with GLIMPSE_phase and the resulting chunks were joined together using GLIMPSE_ligate. Variants were filtered based on genotype likelihood scores > 0.99.

## 3. Results

### 3.1 Dental calculus proteomic analysis

We performed a palaeoproteomic analysis of dental calculus from 77 individuals excavated from 22 archaeological sites, spanning the Pottery Neolithic to the Middle Ages (Figure 2). To evaluate protein preservation and detect potential contamination, both ancient dental calculus samples and laboratory controls were screened against the Oral Signature Screening Database (OSSD; see Methods). Of the 77 analysed samples, 61 (79%) exhibited well-preserved oral signature proteins, indicating their suitability for further analysis (Supplementary Table 2).

**Figure 2:**
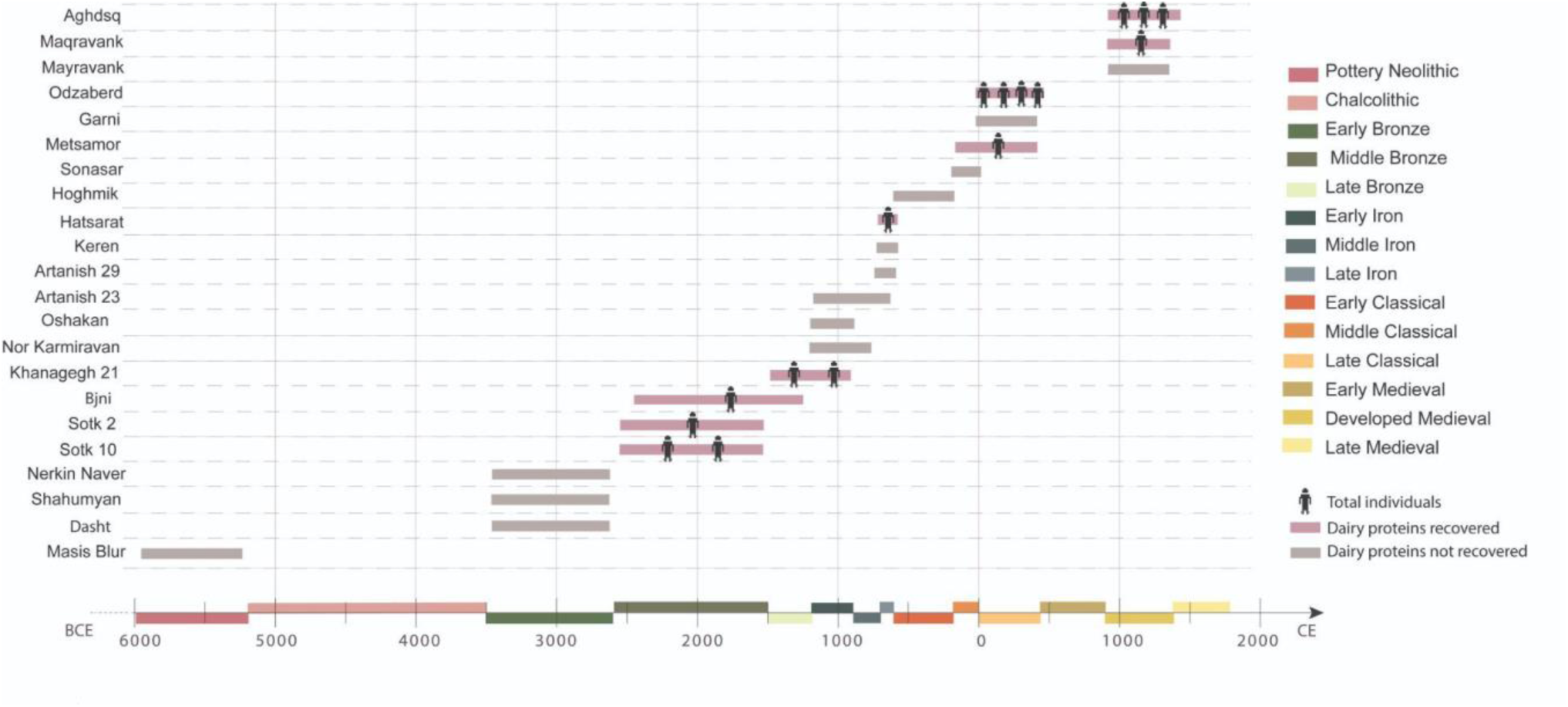
Timeline of sites and individuals analysed through palaeoproteomics

Among the 61 calculus samples that passed OSSD screening, dairy peptides were identified in only 15 individuals from 9 sites. The remaining 46 calculus samples didn’t have dietary proteins. The detected dairy peptides were predominantly derived from β-lactoglobulin (BLG), identified in 13 samples (Figure 3), and β-casein, detected in 2 samples. Most individuals with positive dairy protein results exhibited multiple peptide spectral matches to ruminant dairy proteins (Supplementary Table 3). While some milk peptides were only specific to higher taxonomic levels (Pecora infraorder (n=22 peptides), Bovidae family (n=2), Caprinae subfamily (n=6)), others allowed for more refined taxonomic classification. In particular, peptides were assigned to milk from sheep (Ovis, n=4), goat (Capra, n=7), and/or cattle (Bos/Bovinae, n=4). At the same time, some of the detected milk proteins consisted of bovid peptides shared between both sheep and cattle, suggesting the consumption of milk from either species (Ovis/Bovinae, n=55).

**Figure 3:**
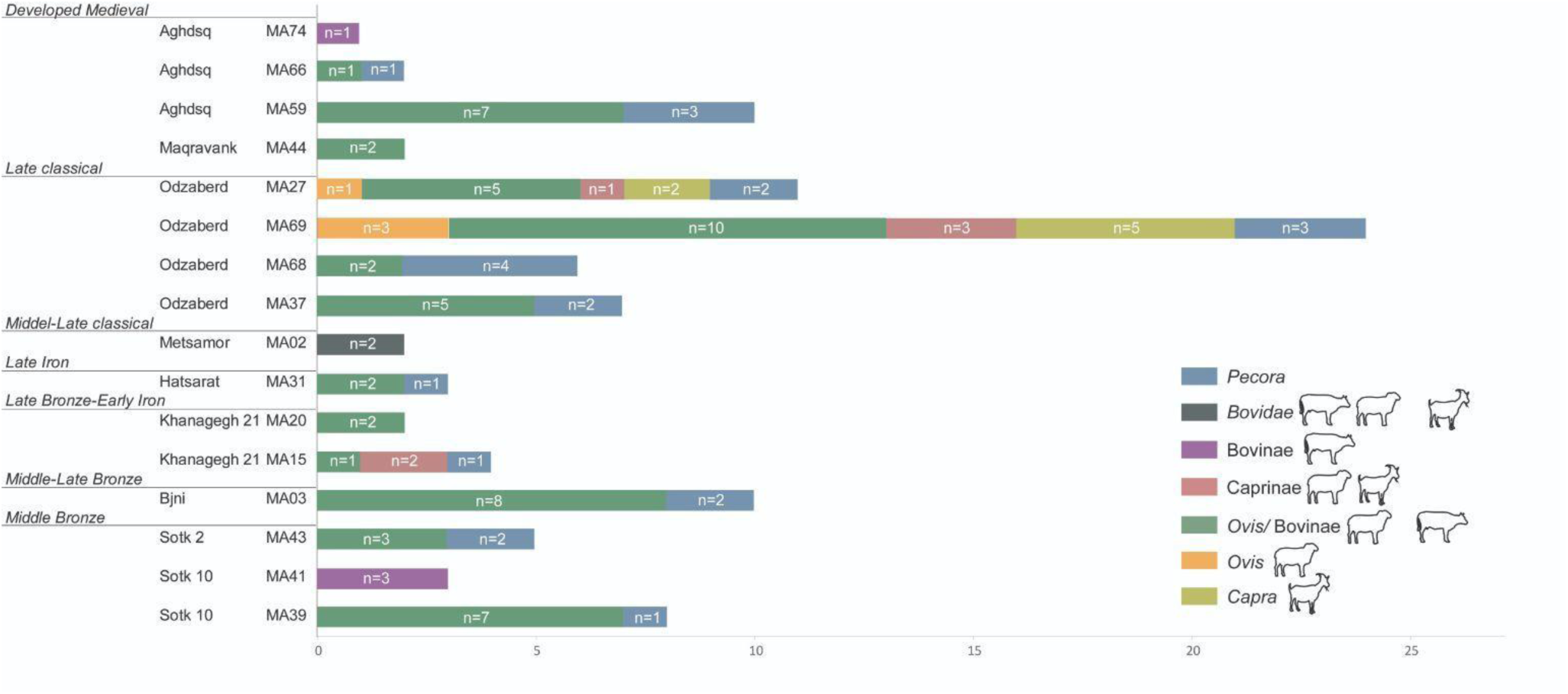
Taxonomic specificity of milk protein spectral matches per individual

The presence and distribution of milk peptides fluctuate across different periods. The earliest individual in our study, MA100, an adult male between the ages of 45 and 60, excavated from the Late Neolithic Masis Blur site, exhibited well-preserved oral proteins but showed no direct evidence of milk consumption.

Similarly, among the nine Early Bronze Age individuals analysed, four displayed well-preserved oral proteomes; yet, none contained detectable dairy proteins. The earliest direct evidence of milk consumption in our dataset comes from the Middle Bronze Age, with all four individuals analysed exhibiting milk proteins. Specifically, individuals MA39, MA43, and MA03, from the Sotk-10 site, preserved β-lactoglobulin peptides specific to Pecora and Ovis/Bovinae, while individual MA41, from the same site, contained peptides specific to cattle.

The Late Bronze–Early Iron Age assemblage includes dental calculus from six individuals, with only one specimen exhibiting poor preservation of the ancient oral proteome. Two individuals from the Kanagegh-2 site yielded milk peptides: MA15 preserved BLG peptides assignable to Pecora, Ovis/Bovinae, and Caprinae, whereas MA20 displayed peptides attributed to Ovis/Bovinae. Among the ten individuals assigned to the Middle to Late Iron Age, six passed OSSD screening, but only one individual (MA31), from the Hatsarat site, exhibited evidence of milk consumption, with BLG peptides specific to Pecora and Ovis/Bovinae.

In the Early to Late Classical period, seven individuals were analysed, of whom only one did not preserve proteins, while one individual (MA02), from the Metsamor site, contained β-casein peptides specific to Bovidae. The Late Classical period is represented by ten individuals, eight of whom exhibited a well-preserved oral proteome. Of these, four individuals from Odzaberd had evidence of milk proteins: a male aged 50-55 (MA37) and a female aged 35–45 (MA68) displayed BLG peptides assignable to Pecora and *Ovis*/Bovinae, while male aged 14-17 (MA69) and male aged 35-45 (MA27) provided direct evidence of milk consumption from multiple taxa, with BLG peptides unique to both sheep and goat, as well as *Ovis*/Bovinae. The Medieval period includes 31 individuals, with only two failing the OSSD screening. Among the remaining 29 individuals, only four contained detectable milk proteins: MA44 exhibited BLG peptides matching Ovis/Bovinae, MA59 and MA66 displayed BLG peptides specific to Pecora and *Ovis*/Bovinae, and MA74 preserved β-casein peptides specific to Bovinae.

### 3.2. Allele distribution in modern and ancient populations in Armenia

We explored the presence of five SNPs associated with LP in modern Armenian populations. In Armenian samples, we found a modest frequency (8%) for the 13910C>T variant (Supplementary Table 4), which is the most commonly observed mutation in European populations. This variant is found in high frequency in India, northwestern Africa, and the western African Sahel, but is generally not prevalent in most East African and Middle Eastern populations. Interestingly, the other four variants (−13915T>G, −14010G>C, −13907C>G, and −14009T>G) were absent in Armenian samples (Figure 4 ; Supplementary Table 4). This pattern is noteworthy given Armenia’s geographical position at the crossroads between Europe and the Middle East, with the latter region having the −13915T>G allele as the most prevalent variant associated with lactase persistence. The 13910C>T locus showed no significant deviation from Hardy-Weinberg equilibrium in the full genome dataset representing the pooled Armenian population, which could be explained by the relative homogeneity of Armenian populations from the different regions within the Armenian highlands.

**Figure 4:**
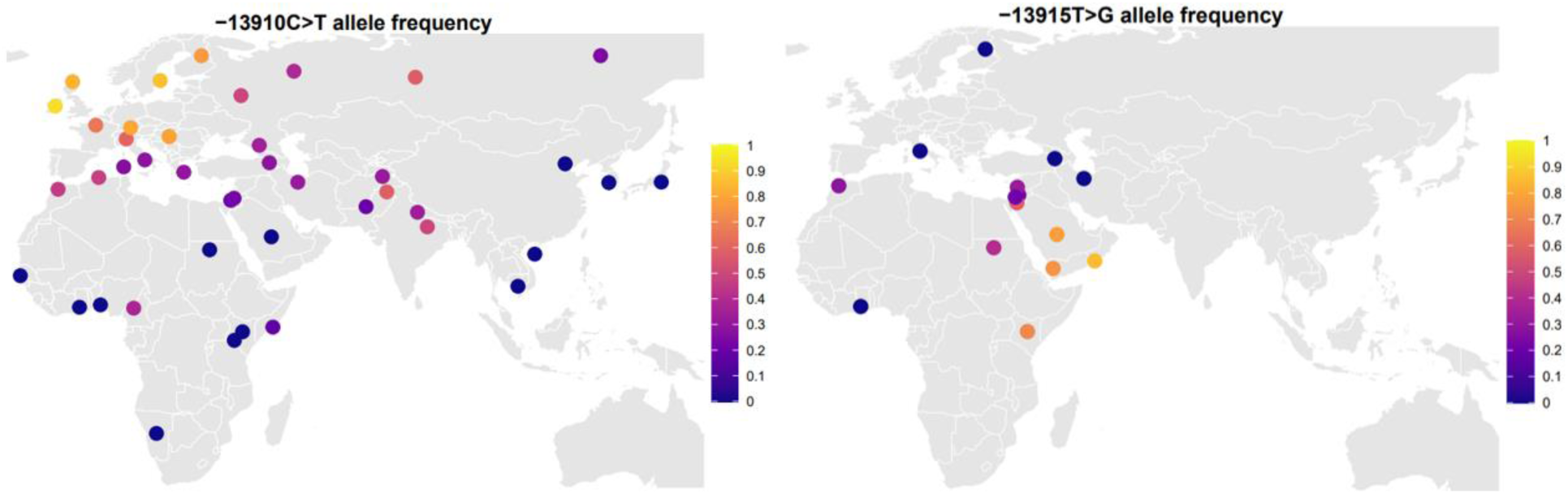
LP variants in modern populations

We next screened for the presence of LP-associated variants in all published ancient samples from Armenia. Following imputation, nine samples showed positive results for the derived allele of rs4988235 (Supplementary Table 5). To confirm these results, we directly screened reads covering this position using stringent filtering criteria. While this position was not covered in most low-coverage ancient samples, we successfully confirmed the presence of the derived allele in a sample from Harzhis, dating to the Late Urartian period (680-550 BCE). The results for the other eight samples, including Early Bronze Age specimens from Armenia (2,600-2,300 BCE), require cautious interpretation, as they could represent false positives from imputation. In this scenario, they could have carried haplotypes which have ancestral rs4988235 genotypes but which are similar to that on which the −13910C>T variant arose (Burger et al. 2020; Ali et al. 2022).

We then aimed to directly check reads for the previously reported oldest case of the derived allele in the sample from Ukraine (I6561), dated to ∼4,000 BCE (Mathieson et al. 2018). With our stringent filtering approach, the sample yielded only a single sequence read with the derived allele, and given the partial-UDG library treatment method, we could not reject the possibility that this could be due to error. We also screened recently reported samples from Patterson et al. 2022 (Patterson et al. 2022), and the oldest sample we confirmed to carry the derived allele was the individual from the Czech Republic (I7953), dated to 3,000-2,000 BCE.

## 4. Discussion

Our palaeoproteomic findings shed light on the dynamics and specificities of human subsistence strategies in Armenia through time, particularly regarding the emergence of milk consumption, its intensification, and diversification. Based on milk proteins identified in the dental calculus of 15 individuals from 9 sites, we demonstrate that milk products were consumed in the region by at least the 2nd millennium BCE (Middle Bronze Age), with cattle and possibly sheep (indicated by non-specific bovid peptides, which could derive from either sheep or cattle) serving as the primary sources of dairy products in these early dairying cultures. We find a continuity of cattle and/or sheep milk consumption in the region from the Middle Bronze Age through to the Medieval period. Over time, both the intensity of milk consumption and the diversity of exploited animals increased, with the first direct evidence of goat milk appearing in the Classical period. Noteworthy, milk proteins in dental calculus were predominantly recovered from high-altitude sites situated above 1,500 m asl. This pattern may reflect milk consumption as an adaptive strategy to cope with the harsh environmental conditions of highland regions. A similar trend was observed in a study of milk residues in ceramic vessels (Manoukian et al. 2022) from the Late Bronze Age, where milk use appeared to be more common in upland areas.

When contextualised with existing regional data (Scott et al. 2022; Manoukian et al. 2022), our results contribute to a more comprehensive understanding of the development and role of dairying in the South Caucasus as a whole. Early evidence for animal husbandry and food-producing economies in the South Caucasus dates to the Pottery Neolithic (6,000–5,500/5,200 BCE), a period characterised by extensive interactions between populations in eastern Anatolia, the South Caucasus, and the Levant (Skourtanioti et al. 2020; Lazaridis et al. 2022). These close contacts may have facilitated the spread of animal husbandry as well as the transmission of dairying knowledge and milk consumption practices. Communities associated with the Neolithic Aratashen–Shulaveri–Shomutepe tradition are thought to have lived in lowland permanent settlements with fully sedentary lifeways, subsisting primarily on agriculture and animal husbandry (Badalyan et al. 2022; Chataigner et al. 2014; Badalyan et al. 2004; Martirosyan-Olshansky et al. 2013). While some Neolithic zooarchaeological assemblages (e.g. Masis Blur, Aratashen) indicate caprine exploitation for secondary products, particularly dairy (Berthon 2014; Bălăşescu et al. 2010), no direct evidence for milk consumption has been documented for this period. This absence may reflect preservation biases or the limitations of small sample sizes, though it is also possible that, despite the full domestication and management of ruminants by the late Neolithic, the practice of dairy consumption was not yet widespread in the region. The first direct evidence of milk consumption in the South Caucasus comes from a Chalcolithic individual at the Alkhantepe site, dated to ca. 3,700 BCE, where cattle milk proteins were identified (Scott et al. 2022). Similarly, the earliest evidence of milk consumption in the North Caucasus comes from the Progress 2 site, dated to 4,338–4,074 BCE, while sheep milk consumption was detected (Scott et al. 2022). These results align with abundant zooarchaeological data from the region, where kill-off profiles and birth seasonality patterns suggest that early Chalcolithic pastoralists deliberately managed their herds to optimise access to milk, meat, and fibre (Antonosyan et al. 2025).

During the Early Bronze Age (3,500/3,350-2,500 BCE), the economic importance of pastoralism increased in the South Caucasus, where a unique cultural tradition, named the Kura-Araxes, grew out of Chalcolithic interactions in this region in the mid-fourth millennium BCE. These were non-hierarchical sedentary communities that practised farming and herding (Kohl 2012; Sagona 2014; Badalyan 2014). None of the Early Bronze Age individuals associated with the Kura-Araxes cultural complex, recovered from three archaeological sites and analysed in our study, yielded milk proteins. This is in line with zooarchaeological records from the region, as kill-off patterns of regional sites reflect that sheep and goat were mainly exploited for meat consumption, with no evidence of any special focus on secondary products (Palumbi 2016; Piro 2008; Badalyan et al. 2015). Nevertheless, the absence of milk proteins is somewhat surprising given lipid residue analyses of Kura-Araxes ceramic vessels from seven Armenian sites, which revealed traces of ruminant milk, though at low levels, alongside a stronger dietary emphasis on meat (Manoukian et al. 2022). While fatty acid profiles cannot identify the specific animal source, zooarchaeological data from the region point to a predominance of cattle and caprines, comprising over 50% of the identified faunal remains (number of identified specimens) (Decaix et al. 2019; Piro 2008; Badalyan et al. 2015; Palumbi 2016; Davoudi et al. 2018). Notably, the use of ceramic vessels for processing dairy products appears to have been more prevalent at highland settlements located above 1,000 meters compared to lowland areas (Manoukian et al. 2022), suggesting differentiated management of animal resources across this cultural province. Although milk consumption clearly intensified from the Early Bronze Age onward, this cultural evolution proceeded without corresponding genetic adaptations, as LP-associated variants did not reach widespread distribution in local populations. While we could not directly confirm their presence in these early periods, we can reasonably infer that the ancestral haplotype on which the 13910C>T variant subsequently arose was already present in the region since at least the Early Bronze Age, albeit at relatively low frequencies.

During the Middle Bronze Age (2,500–1,550 BCE), inhabitants of Kura-Araxes communities appear to have shifted away from settled village life in favour of more mobile ways of living, likely sustained by mobile pastoralism (Hammer 2014; Nugent & Selin Nugent 2020; Lau et al. 2020; Badalyan et al. 2003). This transition was accompanied by significant changes in social organisation, including the earliest clear evidence of social inequality, as reflected in the emergence of large burial kurgans across the region (Smith 2020; Smith 2005; Kohl 2012; Sagona 2017; Birkett-Rees 2012). Our results show a reliance on dairy technology for subsistence for these mobile pastoralists, confirmed at two highland sites along the mountainous shores of Lake Sevan (Sotk 2 and Sotk 10, situated at ca. 2,000 m asl), with cattle milk proteins specifically identified at Sotk-10. In contrast, a previously published analysis of an individual from the Hatsarat site, also located in the Sevan Basin, revealed evidence of sheep milk consumption (Warinner et al. 2014). Meanwhile, proteomic analysis of material from the Middle Bronze Age site of Qızqala, in the Aras River Valley of Nakhchivan, detected the presence of goat milk proteins (Scott et al. 2022). These variations likely reflect ecological adaptations, with environmental conditions appearing to have shaped livestock management strategies. Goats, being more resilient to arid climates and scarce water, were favoured in the dry lowlands of Nakhchivan, while the well-watered, pasture-rich Sevan Basin supported cattle and sheep herding.

After nearly a millennium of more mobile lifeways, communities of the Middle Bronze Age chose to establish more permanent agropastoral settlements in the Late Bronze Age (ca. 1,500–1,150 BCE), but now organised in the first complex territorial polities in the region, administered from hilltop fortresses (Smith 2020; Smith 2005; Lindsay & Greene 2013; Badalyan et al. 2003). In our dataset, evidence of ruminant milk consumption persists from the Late Bronze, aligning with the period associated with the Lchashen-Metsamor cultural tradition. This includes a male individual aged 25– 30, recovered from the high-altitude Kanagegh site (1,970 m asl) that showed proteomic evidence of caprine milk consumption. Additionally, a female individual, aged 30-40, recovered from another high-altitude site, Bjni (1,566 m asl), revealed evidence of sheep or cattle milk consumption.

During the Iron Age, evidence for milk consumption becomes more substantial and geographically diverse. An Early Iron Age individual from the Goytepe site in Azerbaijan revealed the presence of sheep milk proteins (Scott et al. 2022). Notably, although evidence of horse milking has been identified in Early Iron Age remains from the North Caucasus (Scott et al. 2022), associated with pre-Scythian groups repopulating the steppe, no comparable evidence exists for horse milk consumption in the South Caucasus. This absence may reflect a sampling bias; however, it could also indicate that horse milking was a steppe-specific adaptation practised by mobile horse-riding groups. Later in the Iron Age, between the 9th and 7th centuries BCE, the Kingdom of Urartu emerged in the basin of Lake Van and expanded its influence across what is now the South Caucasus. Urartu’s animal economy relied on sheep, goats, and cattle, with horses playing a key role in this militarised society (Göcke & Işik 2014; Mirzoyan & Manaseryan 2003). In our study, ruminant milk was detected in the Late Iron Age Urartian Hatsarat burial (1,990 m asl); however, the milk proteins consisted of non-specific bovid peptides, indicating either sheep or cattle. Significantly, our earliest confirmed presence of the rs4988235 variant in Armenia also dates to approximately 3,000 years ago, corresponding to this Late Urartian period.

In the Late Classical period, when the kingdom of Armenian Arsacids came to power in Armenia, our dataset reveals a notable shift in dairying strategies, marked by the exploitation of a broader range of livestock species. Among eight individuals analysed from the Odzaberd site (1,963 m asl), two exhibited a diversified milk protein profile, including evidence of sheep, goat, and cattle milk consumption. The consumption of milk from multiple species may indicate the wealth or trade connections of the studied individuals; however, in the South Caucasus, mixed herding practices involving the concurrent management of cattle, sheep, and goats were common, reflecting a broader regional strategy of diversified livestock exploitation. Notably, this represents the earliest evidence of goat milk consumption within our assemblage. The first appearance of goat milk in the highlands of Armenia may indicate a diversification in dairy exploitation strategies. However, this pattern could also reflect sampling bias and requires further investigation.

Lastly, we present the only regional dataset documenting milk consumption during the Medieval period. In the Developed Medieval period, our findings suggest a potential correlation between social status and milk consumption. Of the 24 individuals analysed from the Aghtsg burial complex (1,262 m asl), only two showed evidence of sheep or cattle milk proteins. Notably, both individuals had recovered from sarcophagi, indicating their elevated social status. The detection of dairy proteins exclusively in high-status individuals may indicate preferential access to milk products among the elite. These findings underscore the need for further research into the interplay between social hierarchy and subsistence in the Medieval period.

To complement our historical analysis, we also investigated the prevalence of LP alleles in the modern Armenian population and whether the LP-associated variants in the South Caucasus are primarily of European or Middle Eastern origins, thereby addressing a gap in previous studies of LP allele distribution (Anguita-Ruiz et al. 2020). Interestingly, we found ∼8 percent prevalence of the European LP allele and complete absence of the predominating Middle Eastern variant. However, we speculate that unidentified novel LP variants may be present in the population, necessitating GWAS analysis for comprehensive detection. Alternatively, the relatively low frequency of LP-associated variants compared to many other populations may reflect the predominant use of fermented dairy products such as matsun, which contain reduced lactose levels.

Thus, this study presents the first large-scale palaeoproteomic and genetic investigation of milk consumption and genetic adaptation to it in Armenia, extending from prehistory to modern times. Through a comprehensive analysis of preserved dental calculus from individuals across all studied archaeological sites, our dataset captures an unprecedented temporal and geographic scope. While sample size is constrained by material preservation, this approach provides a robust foundation for exploring the emergence, development, and diversification of dairying practices in the South Caucasus, revealing key patterns in how milk consumption evolved across different environmental and cultural contexts. Similarly, genetic analyses of both ancient and modern samples from Armenia offer a comprehensive longitudinal view of lactase persistence development within one geographic region, tracing its emergence and spread through the local population over thousands of years. The study of over 6,000 years of dairy pastoralism in the South Caucasus demonstrates that, despite early and intensive dairying in the region, genetic adaptation through LP arose later and has never become widespread, remaining relatively rare even among contemporary Armenians, even though dairy continues to be important in the diet. These findings underscore that cultural and dietary strategies— such as fermentation, cheese-making, or mixing milk with other foods—rather than genetic adaptation alone, enabled populations to sustain and benefit from dairying. Key questions nonetheless remain regarding the origins of dairying and whether milk consumption served as the primary selective pressure driving the evolution of lactase persistence, or if other factors contributed to this genetic adaptation. Further ancient DNA studies with direct screening evidence are required to shed light on the question of origins and spread of the LP alleles in the region.

## Supporting information

Supplementary textt

## Data availability

LC-MS/MS data files are available at ProteomExchange and were uploaded through MassIVE in two batches. The files can be accessed with an FTP client using the download links: https://ftp://MSV000098912@massive-ftp.ucsd.edu for Batch 1 and https://ftp://MSV000098915@massive-ftp.ucsd.edu for Batch 2.

## Acknowledgments

The research was funded by the Max Planck Society. L.Y., Z.K. and S.M. are supported by the Higher Education and Science Committee, MESCS, Armenia (Projects #21AG-1F025).

