## Supplementary textt for "Palaeoproteomic and genetic insights into millennial-scale dairy consumption in Armenia"

### Archaeological site descriptions

**Aghtsk**: The royal tomb of Aghtsk is located in the Aghtsk community of the Aragatsotn Province of the Republic of Armenia. The monument has been known since as early as the 4th–5th centuries AD in written sources. This multi-layered site was excavated periodically between 2015 and 2022. In the central part of the site, a medieval cemetery was uncovered. So far, the excavated remains are around 256 individuals from the High and Late Middle Ages, of which 185 have been identified (18 newborns, 61 children, 47 females, 43 males, and 16 undetermined).

The burials followed Christian funerary practices—bodies were laid on their backs, generally facing east, with extended torsos, hands crossed over the chest or abdomen, and legs placed parallel. The skeletons were found at depths of 30–100 cm below the surface, densely packed in 3–4 layers, often directly on top of one another. They date to the High and Late Medieval periods (Safaryan, 2021).

**Artanish 23** is an Iron Age necropolis located in the northwestern part of the Artanish Peninsula (1,927 m asl, Gegharkunik Province). It was discovered at the beginning of the 20th century by Yervand Lalayan. The Armenian-German expedition carried out surveys here in 2015-2016. The cemetery consists of more than 30 visible cromlechs and four huge burial mounds encircled by cromlechs. In total, ten tombs were excavated during 2019-2022. Tomb No.1 has a round cromlech; the chamber is oriented east-west and is made of slabs. A 30-35-year-old male was buried in the tomb. Materials were documented in situ, including vessels, bronze and iron arrowheads, bronze bracelets, rings, and various beads. Faunal composition at the tomb is demonstrated only by sheep/goat and cow/bull (Bobokhyan and Kunze, 2021; the osteological material is identified by N. Zarikyan).

**Artanish 29** is an Iron Age necropolis, located in the western part of the Artanish Peninsula (1935 m. asl, Gegharkunik province), ca. 1 km away from Artanish 23. The cemetery consists of more than 20 tombs encircled with round cromlechs. It was discovered by the same expedition in 2015. In 2019 and 2021, two tombs were excavated. Tomb No. 1 is a burial mound encircled with a round cromlech and has stone-soil armour. A cist grave was opened in the centre of the cromlech with the west-east orientation. The walls of the chamber were lined up quite neatly and canonically. Meanwhile, the biological and archaeological materials inside had no canonical arrangement. The chamber was full of human bones, animal bones (sheep/goat, cow/bull, pig, wolf; the osteological material is identified by N. Zarikyan), ceramic sherds, beads, metal, and bone objects that were buried here simultaneously (Bobokhyan and Kunze, 2021).

**Dasht:** The site is located in the Armavir region, on the 8th km of the Ashtarak-Vagharshapat highway, 150 m south of the Dasht village, at an altitude of 947 m above sea level. It is an Early Bronze Age tomb "kurgan" discovered in 2021 by the rescue archaeology team of the Institute of Archaeology and Ethnography NAS RA (field director L. Aghikyan). The cist chamber contained around 27 human individuals, more than 2000 beads, 12 vessels, and 2 bronze objects. All the artifacts belong to the Elar-Aragats type of the first phase of Kura-Araxes culture and date back to the last quarter of the 4th millennium BC (Aghikyan and Badalyan, 2025).

**Hatsarat**: The tomb was uncovered in 2016 during construction work at the Berd Glukh archaeological site, located within the administrative boundaries of the Hatsarat community. The burial chamber is soil-cut and oriented east to west. At a depth of about 1.15 meters, artifacts dating to the late period of the Kingdom of Van (Urartu), specifically the 7th century BCE, were discovered. These included red polished ceramic vessels, two glass beads, a fragment of a bronze object, and a piece of a bone inlay decorated with a carved motif—likely part of an inlaid item belonging to another object.

**Hoghmik:** Hoghmik archaeological site is located 4 km southeast of the town of Amasia, Shirak Province, Armenia, on a rocky promontory on the right bank of the Nili, a tributary of the Akhuyan River, at an elevation of up to 2000 m above sea level. It lies within a pastoral zone, in an environment of predominantly treeless alpine meadows enriched with black soil. The site is dated from the 2nd century BCE to the 4th century AD. The associated material is diverse and includes various local and imported ceramics, stone vessels, ornaments, terracotta figurines, ostraca, osteological and zoological material, and more.

**Kanagegh:** The Kanagegh archaeological site is located on the western shore of Lake Sevan, about 5.5 km north of the village of Yeranos, in the Gegharkunik region of Armenia. The site consists of a settlement and a burial field, occupying approximately 200 hectares. In the burial field, 30 burial mounds have been preserved. From 1980 to 2024, eleven of these were excavated. Archaeological finds indicate that the earliest burials in the cemetery date back to the 15th–14th centuries BCE. In some burial chambers, reburials were performed during the Early Iron Age (12th–9th centuries BCE) and the Classical period (4th–1st centuries BCE). Excavations in the Kanagegh burial field have uncovered bronze daggers, diadems, bracelets, a Mitanni cylinder seal, stone moulds for casting ornaments, and numerous luxury items (beads, earrings, necklaces, rings, brooches, etc.).

**Makravank** is located on the border of Nig and Varazhnunik (Tsaghkunk) provinces of Ayrarat state of historical Mets Hayk, at the foot of the Tsaghkunyats mountain range. The exact etymology of the name "Makravank" is unknown. Mesrop archbishop Smbatyants supposes that, perhaps, it was a monastery of chaste hermits, or for cleansing from leprosy and other diseases. Information about the monastery has been preserved only in late medieval written sources.

The complex is completely built with polished basalt, consists of a cathedral church (beginning of the 13$^{th}$ century) with a central dome, an eastern two-story depositary, of a ruined courtyard (beginning of the 13$^{th}$ century), probably with cuadricolumn construction, and from the church (10–11$^{th}$ cc.) with single-nave construction and double-sloped roof on the southern side. The churches were restored in the 1980s. The medieval cemetery, rich with khachkars and tombstones, is spread around the complex.

The expedition of the Institute of Archeology and Ethnography NAS RA in 2018 carried out excavations in the monastery complex before the works of partial restoration of the courtyard and beforehand of putting the area of the complex. The courtyard and the surroundings of the two churches were excavated. A lot of archeological artefacts (pottery, metal objects, etc.), khachkars, and tombstones were discovered, and the inscriptions of the monastery were supplemented. The report is dedicated to enlightening the history of Makravank and the results of the excavations (Grigoryan A, Ghulyan A, Miridjanyan D, 2023).

**Masis Blur**: Masis Blur lies in the Ararat Valley (Araxes River Basin) at 862 m. above sea level (asl), in Ararat province. The site was first surveyed in 1969, and the mound was subsequently destroyed during the construction of a greenhouse complex. Masis Blur is a late Neolithic Aratashen-Shulaveri-Shomu type settlement dated to 5980-5490 cal BCE. The zooarchaeological evidence from Masis Blur suggests that animal husbandry, particularly sheep and goats, was an important component of the subsistence system. Wild animals (wild aurochs, wild boar, horses, red deer, gazelle, hare, fox, hedgehog, and wild caprines) are also present in very low numbers (Olshansky, 2018).

**Mayravank**: Hovhan Mayravanetsi, who was a prominent public figure, philosopher, and theologian of the 7$^{th}$ century, was one of the most visible persons of the Armenian reality, irreconcilably non-Chalcedonian. In 506, after the church Council in Dvin was held by the Catholicos of All Armenians, the tense relations between the Armenian and Greek churches at the end of the 6$^{th}$ and the beginning of the 7$^{th}$ century reached their peak. In 633, Catholicos Ezr A. Parazhnakertsi accepted the doctrine of monothelitism and established unity with the Byzantine Church in the Armenian-Byzantine Council held in Karin with the participation of the Byzantine Emperor. Relations with the Catholicos became so strained that Hovhan Mayravanetsi, a supporter of the independence of the Armenian Church, was removed from the Dvin Catholicosate and settled in Mayravank.

Mayravank is located 3 km north of the village of Solak, Kotayk region, RA, at the height of one of the wooded gorges of the Tsaghkunyats mountain range (coordinate: N40°29'6.99" E44°41'21.90", altitude: 1980 m asl.), in the Nig region of the Ayrarat province of historical Mets Hayk. Many Armenian and Georgian written sources about the monastery begin and end with the activities of Hovhan Mayravanetsi. There are lithographs preserved on the walls of the St. Astvatsatsin, which fill especially the unknown pages of 13$^{th}$ century.

The report is dedicated to the excavations of 2021 carried out mostly for the restoration works in Mayravank; the artefacts were related to the period of Hovhan Mayravanetsi and the subsequent history of the monastery. Archaeological studies show that the second period of the monastery's flourishing was in the beginning of 13$^{th}$ century, when a central dome St. Astvatsatsin Church was built here, probably on the place of the destroyed old one, two-storied on the eastern side, a pair of sacristies and polished walls, khachkars were erected here. Most of the artefacts unearthed during the excavations in the church and its surroundings also belong to this period: those are pottery shards, metal objects, and stone tools. Anthropological remains from burials were also found during the excavations on the northern side of the St. Astvatsatsin Church (Grigoryan A, Ghulyan A, Simonyan H 2023).

**Metsamor**: The Metsamor archaeological site is situated in the Ararat Valley, 35 km southwest of Yerevan, close to the marshy sources of the Metsamor River. The first information referring to this site was recorded in 1890 (Alishan, 1890), and the first excavations were carried out in 1959–1962 (Barseghyan, 1962). The settlement was occupied from the Early Bronze Age (Kura Araxes culture) to medieval times (Piliposyan, 2014). The faunal assemblage indicates that cattle, sheep, and goat husbandry were practised at the site. Additionally, horse, donkey, and camel remains were present in the faunal material, together with representatives of the local wild game (red deer, gazelle, beaver, and hare; Jakubiak and Bigoraj, 2020).

**Nerkin Naver**: In 2018, the expedition of the Scientific Center for Research of Historical and Cultural Heritage (dir. H. Simonyan) continued excavations at the site of Nerkin Naver, located 35 km west of Yerevan. The site includes a kurgan field, caves, and circular structures built with large boulders. Among them, Kurgan 4 was of particular interest due to its double-layered burial structure.

In the upper layer, beneath a stone-and-earth mound, a 35–45-year-old male bodyguard was buried. Below, on a specially prepared tuff slab and covered with cloth, lay the severed head of the master. Grave goods included two silver earrings, bronze coiled padlocks, quartz glass and Egyptian faience beads, a glass eye-bead (possibly the earliest of its kind), and a decorated clay mug similar to Beden-type ceramics.

The lower layer and undercut northern grave contained scattered multicolored beads (black, white, turquoise, amber), finely worked carnelian, black amber, and gold-plated gypsum beads of exceptional craftsmanship. Other items included red- and black-polished vessels, a bronze dagger, a pin, a mirror-standard, a basalt piala, seashells likely from the Persian Gulf, obsidian flakes, pearl shells, and remnants of matting. Particularly notable were gold-plated gypsum wheels—likely from a model ritual cart intended to transport the soul of the deceased.

Although small (8.5–9 m in diameter, 0.35 m high), the kurgan contained rich and diverse burial goods. Parallels were noted with materials from Beden (Berikldebi) and Martkopi (Shindatkhevi), especially in ceramics, burial customs, and the symbolic role of a bodyguard. Certain ceramic forms also resemble those from Shengavit.

Based on analogies with Beden and Martkopi materials and the presence of red-polished ceramics, Kurgan 4 is dated to the early stage of the Middle Bronze Age, ca. mid-3rd millennium BCE. Microscopic analysis of residue in vessels revealed a porridge made of wheat, eggs, and crab. Chemical and X-ray analyses confirmed that the high-quality beads were made from quartz glass, which requires extremely high temperatures—an impressive technological feat for the time.

The evidence from Kurgan 4 suggests that advanced glass and metalworking technologies were present in the South Caucasus by the mid–3rd millennium BCE, supporting the view that a highly organized and technologically sophisticated society existed in the region, comparable to contemporary cultures of the ancient Near East (Simonyan, 2010)

**Nor Karmiravan**: This site, located in the Martakert region of Artsakh, has yielded significant Iron Age findings. Excavations between 2016 and 2018 uncovered numerous anthropomorphic stelae and various archaeological materials, including slabs and animal bones. These stelae, both clustered and scattered across the site, offer insights into the cultural and spiritual practices of the region's ancient inhabitants. A burial chamber with a cromlech structure, along with the remains of a horse and its bridle, was also discovered. Further precision in dating was achieved through laboratory C14 analysis of one sample from the burial, dating it back to approximately 463 BCE (Yeranyan, Zarikyan, and Simonyan, 2023).

**Odzaberd**: In 2018, excavations were conducted in the southwestern part of Odzaberd's outer city (locus H). In this area, a 16-meter-long section of massive walls built with semi-dressed stones, approximately 2 meters wide, was uncovered. Clay floors, an economic well, and structural fences were also identified, suggesting the remains of a domestic compound enclosed by an outer fence.

Based on preliminary stratigraphic observations, the structure likely dates to the post-Urartian period, specifically the late 7th to 5th centuries BCE. Numerous pottery sherds were found, including fragments of portable hearths, metal objects, animal bones, and other domestic refuse.

At a later stage, most likely during Late Antiquity, burials were made along the southwestern outer corner of the wall, extending northward. These included a collective burial of five individuals and a single jar burial.

The collective burial contained the remains of one child (5–6 years old), two females (14–17 and 35–45 years old), and two males (30–35 and 50–55 years old). The individuals were buried canonically—in flexed positions on their right or left sides—except for one individual whose skeletal remains were disarticulated and found scattered within the burial area.

To the north, a single jar burial was unearthed. Based on the poorly preserved skeletal remains, the individual was a male (35–45 years old), buried in a flexed, embryonic position inside the jar. No accompanying grave goods were found in this burial.

The presence of these burials indicates that by the Late Antique period, the outer city of Odzaberd had been abandoned and was partially repurposed as a burial ground. Anthropological samples were taken from all six individuals and sent for ancient DNA analysis.

**Shahumyan:** The site is located in the Armavir region, around 500 m south of Shahumyan village, at an altitude of 941 m above sea level. The tomb was discovered and excavated by the team of the Institute of Archaeology and Ethnography in 2021 (field director R. Badalyan). The slab chamber contained more than 50 individuals, around 60 vessels, more than 200 bronze objects, and 4000 beads. The materials belong to the Shresh-mokhrablur phase of Kura-Araxes culture and date back to the first half of the 3rd millennium BC.

**Sonasar**: The mausoleum was discovered in the elevated valley of the Sodk region, approximately 10 km from the village of Sonasar, in the historical province of Siunik. The burial dates to the late 1st century BCE. The tomb structure, along with the associated burial and grave goods, was found largely intact. Among the finds were fragments of burial furniture, including carved and decorated legs, as well as personal ornaments and luxury items made of gold, silver, and bronze.

The burial likely belonged to a woman of middle social standing, interred with rich and aesthetically refined objects. Notable finds include a gold necklace with an oval medallion containing green glass, set into a shield-shaped plaque; another necklace with bird-head terminals and a convex agate gem featuring the relief of a vulture; and two gold rings—one bearing an image of Athena standing with an owl, the other with the figure of a standing youth. A pair of earrings depicting a worship scene of the Mother of Gods, Cybele, and her consort Attis, was also recovered.

These items show strong parallels with objects from the tomb of Volta Finta in the Sisian region of Siunik. In particular, a silver cup from that tomb bears an Aramaic inscription naming Araqszat, meaning "protected by the gods"—a title typically reserved for rulers or high officials. This suggests a shared cultural context between the Sonasar and Sisian tombs. Both mausoleums date to the late 1st century BCE, reflecting the wealth, artistic expression, and funerary customs of Siunik's elite class during this period.

**Sotk 2** is a multi-layered site located in the eastern part of Sotk village (Gegharkunik province) at an altitude of 2101 m. asl. The earliest cultural layer described at the site dates back to the Early Bronze Age. It is represented by clay architecture specific to the Kura-Araxes culture settlement, together with typical black-polished, but undecorated pottery. The following cultural layer corresponds to the end of the Middle Bronze and the beginning of the Late Bronze Age. The site consists of a fortified settlement with architecture made of clay and stone. Rock-hewn pits of varying diameters and depths are unique, serving primarily an economic function. This layer is rich with archaeological materials, indicating that the population of the region was actively engaged in agriculture, animal husbandry, and hunting. The main focus of agriculture was the cultivation of cereals: bread wheat (Triticum aestivum), barley (Hordeum vulgare), and emmer (Triticum dicoccum). The pastoral economy was based mostly on sheep/goat (Ovis aries/ Capra hircus) husbandry. The osteological material also includes bovine (Bos taurus), pig (Sus scrofa domestica), horse (Equus caballus), and red deer (Cervus elaphus). The settlement was inactive around the middle of the Late Bronze Age. The site solely represents a rich intramural burial from the Late Bronze Age: a 20-25-year-old male was buried. The burial included a variety of ceramic, metal, shell, stone, and bone objects. The settlement was last active in the Iron Age, between the XII and IX centuries BC (Kunze et al., 2013). Four of the studied artefacts were excavated from the Middle Bronze Age and another four from the Late Bronze Age layers of the site.

**Vardenik- Azat gyux**: In August 2008, the archaeological team of the "Erebuni" Historical and Archaeological Museum-Reserve (expedition head A. Meshinyan, archaeologist A. Zakyan), initiated conservation excavations at the semi-ruined church in the village of Azat (formerly Aghkilisa), located on the southeastern shore of Lake Sevan, in the Vardenis subregion of Gegharkunik Province. The project was carried out at the suggestion of the Monuments Protection Agency of the RA Ministry of Culture, with the direct support of the "Land and Culture" charitable NGO.

During the period of Azerbaijani habitation, the village and its Christian monuments had suffered significant damage. Therefore, the systematic excavation and cleaning of the site were deemed an urgent and necessary undertaking.
